# SCAMPI: A scalable and accessible micropatterning platform for reproducible organoid morphogenesis

**DOI:** 10.64898/2025.12.29.696843

**Authors:** Federico Cantoni, Matteo Fiori, Lorenzo Mattolini, Dries Janssen, Elena Ramis-Bravo, Francesca Sgualdino, Julie Van Lent, Jeroen Lammertyn, Adrian Ranga

## Abstract

Micropatterning technologies enable biomimetic organoid generation with controlled in vitro morphogenesis but current approaches lack scalability. Here, we develop a mask-based photopatterning platform to enhance organoid culture reproducibility and scalability on diverse substrate formats. We demonstrate its use in generating architecturally defined neural organoids of varying geometries with reproducible morphogenesis. This technology provide a versatile and low-cost strategy for generating geometrically controlled organoids.

## Main

Organoids recapitulate key aspects of human tissue and organ formation in vitro^1^, however, the stem cell aggregation approaches which are conventionally used to generate them lead to poorly reproducible tissue architectures and variable cell fate patterns, limiting both mechanistic insights and translational applications^2,3^. Micropatterning has emerged as an alternative approach to initiate organoid formation: starting from cells on a micropattern enables precise spatial control over initial cellular organization^4^, leading to more defined 3D morphogenesis. Several groups have developed methods to control multicellular shape and identity using micropatterning technologies in a variety of model systems. Human pluripotent stem cells (hPSC) seeded on circular adhesive islands were used to understand the role of geometry in early cell fate decisions, identifying the critical length scales and morphogens required for the emergence of germ layer patterning^5^. The combination of micropatterned pluripotent stem cells with morphogen gradients used for differentiation has been shown to bias early fate specification^6^ and to generate neural-tube regionalization^7^. Intestinal organoids have also been generated with defined shape, size, and cell distributions, thereby producing predictable and reproducible structures^2^. In the context of morphogenesis, geometric constraints by specifying micropattern size have been shown to be critical in enabling the emergence of neural-tube-like organoids which recapitulate the bending, folding and closing of the neural tube^8^.

In these studies, microcontact printing and single photon light-assisted patterning have been the most widely techniques to generate micropatterns. Microcontact printing uses PDMS stamps fabricated from lithographed masters to transfer proteins onto substrates, to which cells then adhere. However, this approach requires contaminant-free (eg. particles and fibers) stamping surfaces, uniform contact pressure and is unsuitable for nested geometries. Additionally, this technique necessitates mechanical supports for features smaller than ∼100 μm due to stamp instability^9^, and is impractical for large surfaces or for high-throughput well plates.

Conversely, UV light-assisted patterning enables the patterning of features with higher resolution, as well as the potential for higher scalability^10^, both in terms of feature sizes (minimal feature size ∼ 1µm ) as well as in throughput. Nonetheless most reported implementations of this technique have relied on digital-micromirror–based systems, which allow for rapid pattern design but are expensive, require specialized equipment, and necessitate time-consuming sample rastering due to limited field-of-view. To address these limitations, we developed a **SC**alable and **A**ccessible **M**icro**P**attern**I**ng platform for organoid morphogenesis (**SCAMPI**). This system is an affordable (<€1000), standalone, and portable device based on mask patterning which uses previously developed PEG-coated substrate chemistries^11^ to enhance accessibility and scalability for organoid cultures with defined geometries (**Fig.1**).

**Figure 1.**
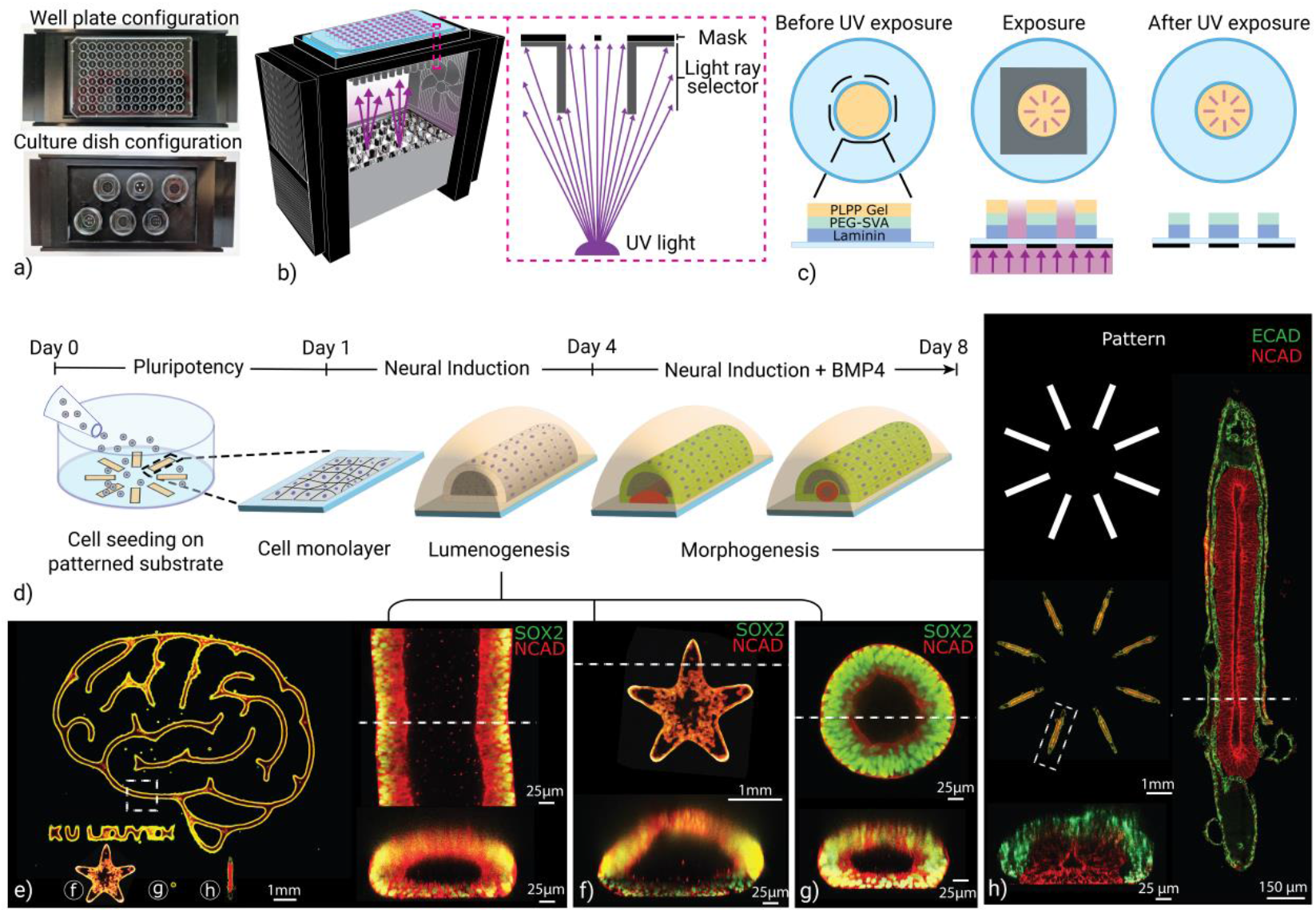
a) System configuration for well plates and glass-bottom cell culture dishes. b) Schematic of the SCAMPI device and inset of the light ray selector design. c) Steps of substrate PEG passivation and patterning with UV light. d) Experimental scheme showing the steps to obtain hPSC-derived neural organoids. e-g) Single-lumen neural organoids obtained starting from patterns generated by the SCAMPI device (day 4), with their respective cross-sections, highlighting the variety of organoid shapes and sizes possible with the device. h) Designed pattern and generated pattern of double-lumen neural organoids at day 8, with magnification of one organoid in the array and its cross-section.

The system employs an inexpensive UV-LED array to transfer a mask pattern to a photosensitive layer on substrates ranging from conventional culture chambers to microwell plates (**Fig.1a-c, Sup.Fig.1a**). Here, the mask is placed in contact with the glass or plastic bottom of the substrate, which eliminates direct contact between the mask and the sample to ensure sterility and throughput without the need for cleaning steps between UV light exposures. A critical element with mask-based system which are not in direct contact with the photosensitive material is the need to minimize light ray divergence in order to limit lateral patterning error (**Fig.1b, Sup. Fig.1b, d and e, Sup. Fig. 3)**. In our system, we implemented a 3D printed light ray angle selector to filter rays exceeding a defined angular thresholds. The implementation of the ray selector results in an energy reduction of 97% and 87% for glass-bottom culture dish and well plate configurations, respectively (**Sup. Fig.1f**). To ensure cost-efficiency and design flexibility both the light ray selector and UV-LED housing were fabricated from black Polylactic Acid (PLA) via fused deposition 3D printing. Black PLA minimizes UV reflection and scattering while offering excellent design flexibility, processability, and smooth surface finishing. The modular housing features snap-fit connections to secure various components of the device and to ensure rapid assembly, maintenance, and substrate exchange within seconds (**Sup. Fig.1 a, b and e**).

To validate the platform for biological applications, we generated human neural tube organoids from hPSCs using neurulation protocols leading to either single-(**Sup. Fig.2a**) or double-lumen (**Fig.1d**) neural organoids which recapitulates the folding and closure morphogenesis of the early neural tube^8^. To induce neural tube morphogenesis, we first maintained the micropatterned 2D stem cell cultures in pluripotency media for a day, followed by treatment with neural induction medium containing TGF-β inhibitor between day 1 and day 8 for single lumen organoids, with the addition of BMP4 to the neural induction medium between day 4 and day 8 for double-lumen organoids . The generated organoids exhibited self-organized lumenization (**Fig.1e-g**) and folding morphogenesis (**Fig.1h**), featuring tubular NCAD+ neuroepithelial tissue, overlayed with ECAD+ surface ectoderm that recapitulated embryonic neural tube cytoarchitecture. Notably, hinge point formation characteristic of neural folding appeared in patterned organoids, demonstrating biomimetic recapitulation of this morphogenetic hallmark (**Fig.1h**). The system demonstrated robust organoid retention on substrates throughout neural tube morphogenesis, with over 90% of seeded patterns successfully maintaining organoid adhesion from initial cell seeding through to mature organoid stages.

SCAMPI enables design flexibility, accommodating µm-to cm-scale features and a variety of geometries which can enable the systematic investigation of the role of geometric variables on tissue architecture and cell fate specification. This versatility extends to the generation of organoids with nested biological features (**Fig.1e**), while maintaining the capability to pattern multiple substrates simultaneously, thereby facilitating high-throughput experimental strategies. To demonstrate these aspects, we used SCAMPI to generate small (150 µm -scale) to large (multi-mm scale) patterns of geometries such as circles, stars or more complex line art. In all of these different pattern geometries, lumen formation was seen at day 4 (**Fig.1e-g**). Interestingly, organoids with widths exceeding approximately 300 µm developed additional walls in the luminal space (**Fig.1f**). Immunofluorescence analysis confirmed the expression of SOX2 and NCAD, indicating successful neural lineage commitment and establishment of neuroepithelial progenitor populations consistent with early neural development. Morphological analysis revealed a continuous single lumen, demonstrating that spatial patterning guided self-organization into architecturally defined neural tissue with structural polarity across different pattern designs **(Fig.1e-g)**.

To verify the reproducibility of SCAMPI, we quantified various biological features generated using the platform. We effectively generated patterns for cell culture, with cells forming a uniform monolayer after seeding (**Fig. 2b, c and e**). A critical capability for system reliability is uniform patterning of large surfaces and different geometries across samples to ensure consistent cell seeding and organoid growth, thereby limiting organoid variability. A small standard deviation of less than 70 cells per organoid was observed after seeding, representing ∼8% of the total cell number per organoid (**Fig.2e**). Immediately following seeding, coverage exceeded the theoretical patterned area but approached expected dimensions after 24 h (**Fig.2f**). This phenomenon was attributed to partially collimated illumination, the UV overexposure to compensate for LED-to-LED power variability and the thickness of the photoactivated hydrogel layer used to remove the PEG-based cell-repulsive coating. This process creates partially exposed zones that extend beyond the theoretical width, where cells initially attach before either spontaneously detaching overnight or migrating toward the central region of the pattern. This geometrical offset was confirmed by laminin staining of the substrate, which showed an average offset inferior to 30 µm for circular micropattern geometries ranging from 150 to 350 µm in diameter (**Fig.2a and d**). After day 1, organoids exhibited growth up to day 5, as well as BMP4-induced morphological changes after day 4, such as flattening of the organoid lumen (**Fig. 2c and f**). By day 8, neuroepithelial cells reproducibly underwent apical constriction, folded and closed, and were enclosed within an outer non-neural ectoderm (**Fig.1h**). This architecture enables investigation of neural tube closure and highlights the system’s capability to replicate in vivo-like morphogenesis mechanisms through organoids, providing a scalable and reproducible platform for modeling early human central nervous system development.

**Figure 2.**
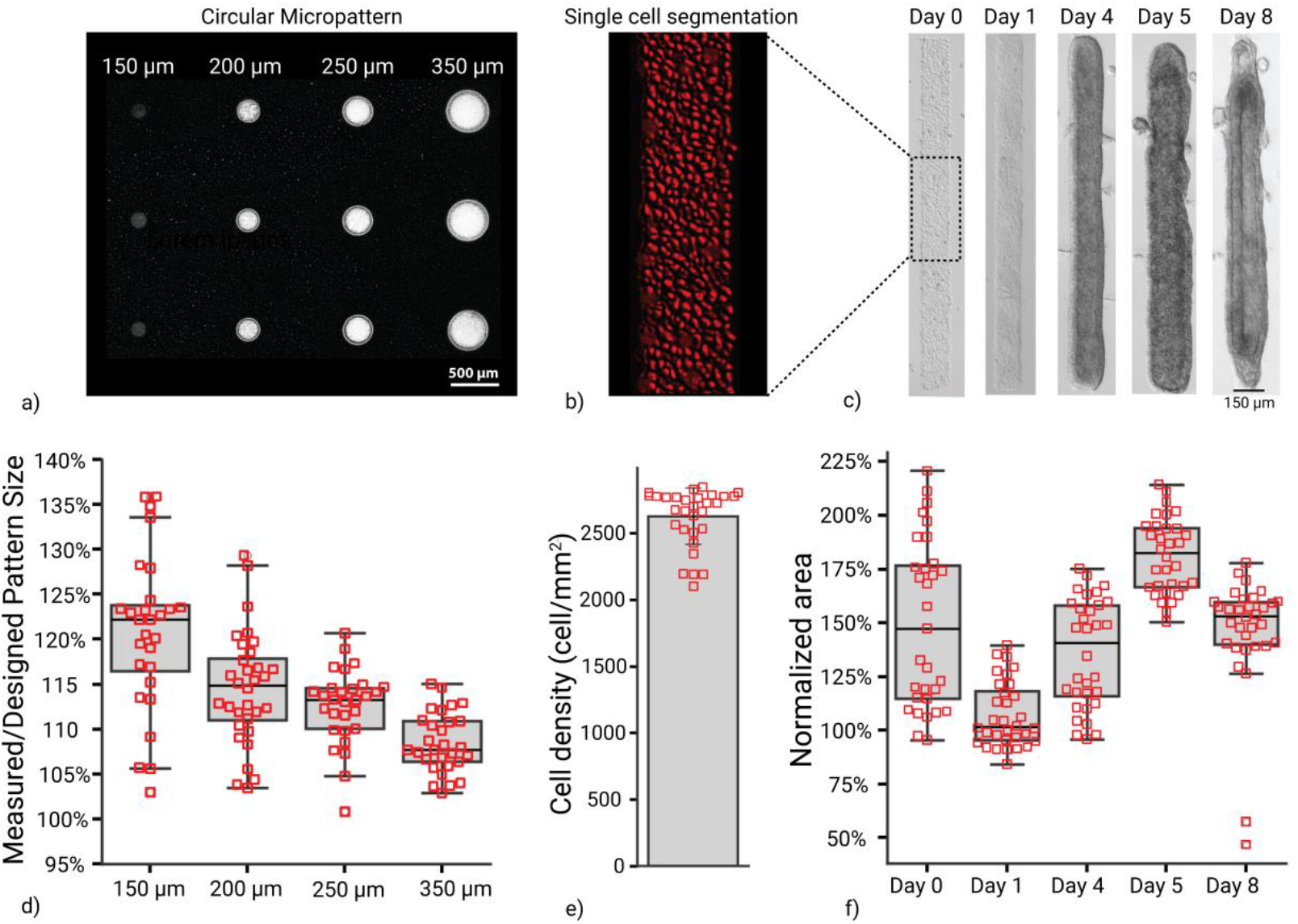
a) Laminin staining of the patterned substrate after UV exposure. b) Magnification of single cell segmentation on micropattern 2 h after seeding (from c), segmented using Ilastik. c) Representative images of double lumen organoid development over 8 days of culture. d) Quantification of pattern fidelity from laminin staining (n=25, 3 independent replicates). e) Quantification of cell density 2 h after cell seeding. f) Organoid area over 8 days of culture (n=31, 3 independent replicates) .

To demonstrate the versatility of the platform in supporting different organoid generation strategies and validate its applicability to alternative differentiation protocols, we exposed cells to the same neural induction medium, including TGF-β inhibitor and 4% Matrigel to obtain neural fate and lumenogenesis, but without the addition of BMP4 between day 4 and day 8. Morphological analysis revealed a continuous single lumen with cells expressing SOX2 and NCAD, demonstrating that also for this model, spatial patterning could guide self-organization into lumenized neural organoids (**Sup. Fig.2b and c**).

Achieving this reproducibility required the implementation of key engineered features for consistent UV exposure and thermal stability. A high-power UV-LED array was selected to minimize patterning time, however this light source also generated elevated temperatures, necessitating active cooling. The 3D-printed device incorporates cooling for heat dissipation during sequential micropatterning cycles while containing UV exposure to a compact and closed footprint (**Sup. Fig.1c and d**). Temperature monitoring showed rapid increase upon activation, plateauing at ∼35 °C within 8 min. Subsequent cycles maintained consistent thermal behavior without heat accumulation, remaining below the 40 °C threshold to prevent substrate softening and device misalignment (**Sup. Fig.1g**). Light intensity monitoring over 10-min exposures with 3-min pauses revealed ∼10% variation, attributed to temperature-dependent LED performance and consistency across cycles (**Sup. Fig.1h**). In addition, LED-to-LED power variance of ∼13% was measured. For the culture chamber, 8-min exposure was chosen to provide adequate margin for inter-LED variability and temporal light intensity decay.

Overall, we show that SCAMPI is a cost-effective and scalable micropatterning platform to generate high-fidelity organoids. The device demonstrates robust performance enabling reproducible sequential patterning cycles. Biological validation using human neural tube organoids confirms the platform’s capability to spatially guide self-organization and morphogenesis, generating architecturally defined tissues that recapitulate key embryonic features including lumenization, biological pattern formation and morphogenesis. The combination of affordability, portability, and high-throughput positions this system as an accessible tool for advancing organoid research and for accelerating translational applications in disease modeling and regenerative medicine.

## Materials and methods

### Design and fabrication of the SCAMPI system

The UV-LED array housing and light ray selector were fabricated via fused deposition modeling (Prusa MK4) using black PLA filament to ensure UV light absorption and to minimize reflection artifacts (**Sup. Fig.1b**). Only the mask holder for the 96 well plate was manufactured via laser cutting (Trotec, 100 Speedy R) a black 3 mm PMMA sheet.

As the UV light of the LED array was not collimated, to determine the UV light incidence angle, an acceptable patterning offset below 15 µm was defined for the 96 well plate substrate. Considering a substrate bottom thickness of 170 µm, an incident angle of ∼ 2.21° was calculated using:

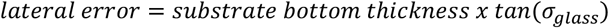

Where *lateral error* represents half the acceptable pattern offset, *glass thickness* denotes the substrate bottom thickness and *σ* indicates the UV incident angle (**Sup. Fig.3a**). According to Snell’s law, an incident angle of ∼ 2.21° in glass corresponds to ∼ 3.37° in air. This calculation assumes perfect contact between the mask and substrate surface, with the polyester mask’s refractive index approximated as equal to that of glass.

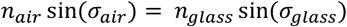

Where *n*_*air*_ and *n*_*glass*_ denote the refractive index of air and glass, respectively, σ_air_ and σ_glass_ indicates the incident light angle of of air and glass, respectively.

For multi-well plates, individual LEDs could not be aligned to each well; therefore, each light ray selector aperture receives light from multiple LEDs. To accommodate pattern sizes up to 4 mm per well (enabling media exchange and cell washing after seeding), a distance of ∼200 mm between the UV light source and mask was calculated to maintain errors below 15 µm, with a light ray selector height of ∼85 mm preventing light from non-adjacent UV-LEDs from passing through mask apertures at angles exceeding 3.37°. The relationship employed was:

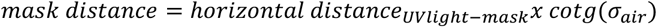

Where *mask distance* represents the vertical distance between the UV light source and the mask, *lateral displacement* represents the horizontal distance between the light source and mask and *light ray* indicates the UV incident angle (**Sup. Fig.3b and c)**.

For glass-bottomed chambers, individual apertures were aligned with single UV-LEDs, with a light ray selector height of ∼75 mm. To ensure adequate heat dissipation from the UV-LEDs, the ray selector was positioned at ∼80 mm, yielding an estimated lateral error below 10 µm (**Sup. Fig.3d)**. Both ray selector designs featured interchangeable mask holders that could be changed within seconds. After the pattern were designed, the flexible masks were printed (Microlitho, UK) and glued to the mask holder using superglue (Cebeo, 175170226) and cured overnight.

### Temperature and light intensity measurements

Light intensity measurements were obtained by aligning the detector sensor with the light collector aperture. Five replicate measurements were acquired at each of six designated positions on the glass-bottom cell culture dish light collector, and at six randomly selected locations for both the UV light source and the 96-well plate light collector. Temporal intensity was characterized using the glass-bottom cell culture dish substrate configuration with its corresponding light ray selector. UV irradiance was monitored over 10-min periods at 20-second intervals for three consecutive cycles during UV-LED array operation. No measurements were conducted during system cooling phases when the UV source was deactivated.

Temperature dynamics were monitored using three temperature sensors positioned at the lowest aperture of the 3D-printed housing. Data acquisition was performed via Arduino microcontroller (Arduino UNO) at temporal intervals matching those employed for UV energy detection.

### Substrate preparation

All experiments utilized 35 mm diameter glass-bottomed culture dishes with 14 mm diameter glass inserts (IBL Baustoff + Labor GmbH). Glass surfaces were activated by 6-min air plasma (Diener electronic GmbH). Subsequently, 120 µL of 0.1 mg/mL of mouse laminin (Becton Dickinson) in PBS (GIBCO) was applied to the inner well and incubated for 30 min at ambient temperature. Following three rinses with HEPES buffer (0.1 M, pH ∼8.4), substrates were passivated by incubation with 0.1 mg/mL PEG-SVA (Hello Bio) in HEPES buffer for 1 h. After four washes with ultrapure water, samples were air-dried at room temperature for 30 min. A photoactivated gel solution, prepared by mixing 2 µL PLPP (Alvéole) gel with 20 µL absolute ethanol (CHEMSOLUTE, 2246.5000), was carefully applied to the inner region of the glass-bottomed culture dish and allowed to dry for 1 h. Photopatterning was performed using the SCAMPI system with 8-min UV exposure to ablate the PEG-SVA layer with the culture dish configuration. Post-exposure, residual PLPP gel was removed by four successive washes with DMEM. To promote cell adhesion within patterned regions, substrates were incubated overnight with 1% Matrigel solution (Discovery Labware). The following day, dishes were rinsed with DMEM to eliminate Matrigel and utilized within 1–7 days.

### Human PSC cell lines and maintenance

All experiments were performed with the KOLF C1 human pluripotent stem cell line (The Jackson Laboratory) at passages below 40. Cells were routinely passaged at 60–70% confluency using 3-min treatment with Gentle Cell Dissociation Reagent (Stem Cell Technology, 100-0485) followed by one PBS wash. Cells were resuspended in 1 mL E8-Flex medium (Thermo Scientific, A2858501) supplemented with 10 µM Rho-associated protein kinase (ROCK) inhibitor (HelloBio, 1:1,000 dilution), then mechanically dissociated by scraping and gentle agitation to fragment colonies. Cell suspensions were split at 1:8 ratio and seeded onto 1% Matrigel-coated 6-well plates in 1.5 mL E8-Flex medium containing 10 µM ROCK inhibitor. Twenty-four hours post-passage, medium was exchanged with 3 mL E8-Flex medium without ROCK inhibitor supplementation.

### Organoid culture on the micropatterns

Double-lumen organoid culture was performed following a previously established protocol^12^, while single-lumen organoids were generated using an adapted protocol. Human PSCs were dissociated from tissue culture plates by 5 min incubation with TrypLE Express (GIBCO, 10043382), followed by enzymatic inactivation using a two-fold volume of DMEM-12 with HEPES (GIBCO). Cells were subsequently dissociated into single-cell suspensions at densities ranging from 7×10^5^ to 9×10^5^ cells/mL in E8-Flex medium (Thermo Scientific) supplemented with 10 µM ROCK inhibitor. A volume of 150 µL cell suspension was seeded onto micropatterned substrates and incubated for 15 min to facilitate cell adhesion. Subsequently, substrates were subjected to three washes with 1.5 mL E8-Flex medium containing 10 µM ROCK inhibitor, with gentle swirling to detach non-adherent cells outside the patterned regions. Samples were then incubated overnight in pluripotency medium (E8-Flex+10 µM ROCK inhibitor), during which residual cells outside patterns spontaneously detached. On day 1, substrates were washed with DMEM to remove detached cells, and 1.5 mL neural induction medium containing 4% (v/v) Matrigel and supplemented with 5 µM TGF-β inhibitor, SB-431542 (Stemgent, 5 μM ), was added. This culture condition was maintained for 48 h to enable morphological transition from two-dimensional cell monolayers to lumenized organoid structures. At day 3, organoids were incubated with neural induction medium supplemented with 5 µM SB-431542. For single-lumen organoid cultures (days 4–8), neural induction medium was exchanged daily. For double-lumen organoid cultures, neural induction medium was supplemented with both 5 ng/mL BMP4 (Peprotech) and 5 µM SB-431542.

### Immunohistochemistry

All organoid samples were fixed with 4% paraformaldehyde, PFA (Sigma–Aldrich) for 1 h followed by 3 washes with PBS. The permeabilization and blocking solution was made with 0.3% Triton X (PanREAC AppliChem) and 3% BSA solution (Sigma–Aldrich) and added to the samples overnight at 4 °C. The following day, primary antibodies were suspended in permeabilization and blocking solution and applied to the fixed and permeabilized organoid for 24 h. Primary antibodies used were: N-cadherin (rabbit, Abcam, ab18203, 1:200), E-cadherin (mouse, Abcam, ab76055, 1:200), SOX2 (goat, R&D Systems, AF2018-SP, 1:200). Three PBS washes were then performed and the samples were then stored for 24 h at 4 °C. Samples were then incubated with secondary antibodies: Donkey anti-Rabbit Alexa 647 (Thermo Fisher scientific, A32795, 1:500), Donkey anti-Mouse Alexa 555 (Thermo Fisher scientific, A-31570, 1:500), Hoechst 33258 (Millipore Sigma, 94403, 1:1000) and Phalloidin-iFluor 488 (Abcam, ab176753, 1:1000) in the permeabilization and blocking solution overnight at 4 °C. Samples were washed 3 times in PBS and imaged. For the tube in a tube organoids, the samples were cleared for 1 h at room temperature in RapiClear 1.47 (Sunjin Lab). For the laminin staining of the patterns, 4% PFA was used for 1 h followed by 3 washes with PBS at 5-min intervals. The samples were then blocked with 3% BSA in PBS for 30 min and then incubated in primary antibody: Laminin (rabbit, Thermo Fisher scientific, PA1-16730, 1:200) at room temperature. After 3 washes with PBS at 5-min intervals, the samples were incubated in secondary antibody: Donkey anti-Mouse Alexa 647 (Thermo Fisher scientific, A-31571, 1:500) for 30 min at room temperature. Finally, samples were washed 3 times in PBS and imaged.

### Imaging and image analysis

Brightfield images were taken using a Zeiss Axio Observer Z1 inverted microscope with a Colibri LED light source and a 10X air objective, and an inverted EVOS microscope using 4X and 10X air objectives. Fluorescence imaging of whole-mount immunostained samples and live-cell imaging was carried out using a Leica SP8 confocal microscope with either 10X or 25X objectives. Both fluorescence and brightfield images were taken directly on the glass bottom of the dishes in which the organoids were cultured. Live imaging was carried out at 37 °C temperature and 5% CO2. Image segmentation of the single cells after seeding (**Fig.2.b**), organoids and the laminin-stained patterns was performed with Ilastik (1.4.1rc2)^13^. The obtained segmented images were then analyzed with Fiji^14^.

## Code availability

The STL files for the UV-LED array case and the light ray selector used in this manuscript are deposited at GitHub: https://github.com/fedekanto/SCAMPI-project.git

## Acknowledgements

This work was supported by FWO PhD fellowships 11K5722N (L.M.), 1S43421N (J.V.L.) and 11M5323N (F.S.), FWO grants G086622N and G0ACA24N (A.R.) KU Leuven grants C14/22/108, C32/17/027 (A.R.) and IDN/20/007 (A.R.), and by an Allen Distinguished Investigator Award (A.R.), a Paul G. Allen Frontiers Group advised grant of the Paul G. Allen Family Foundation. We acknowledge Fablab Leuven for providing access to the facility and manufacturing support, and Thomas Pilkington and Marc Lambaerts for support with the 3D printing.

## Supplementary Figures

**Supplementary Figure 1.**
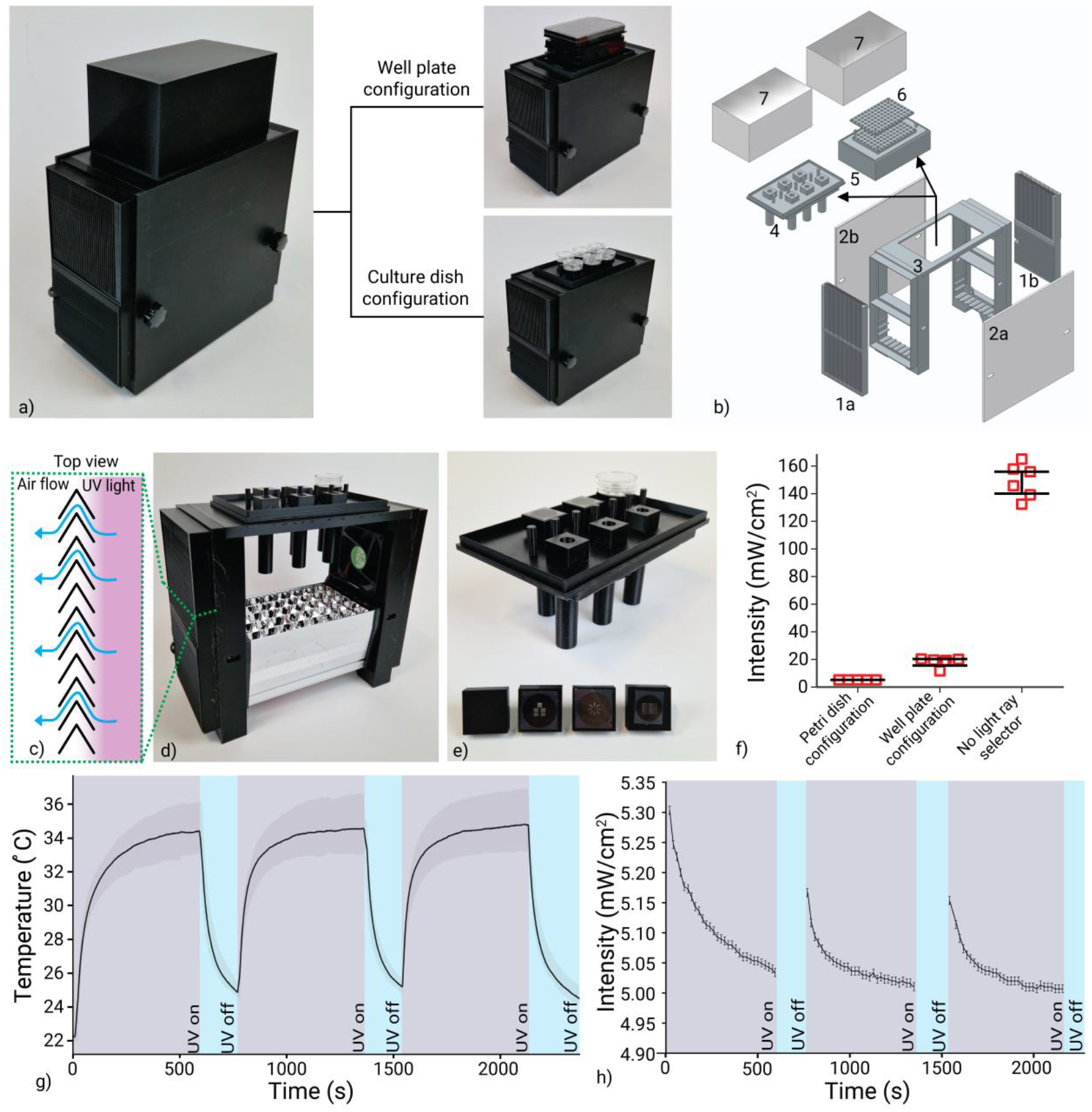
a) Complete system and the two ray selector configurations for well plates and culture dishes. **b)** Schematic of 3D-printed system components: (1) heat exchanger preventing UV diffusion outside the system, (2) side cover, (3) main structural support for component attachment, (4) ray selector configuration for culture dishes, (5) ray selector configuration for well plates, (6) Mask holder and (7) lid. **c)** Top view of the heat exchanger design allowing airflow while preventing UV light diffusion. **d)** Side view of the system without lid and side covers. **e)** Culture dish ray selector with four interchangeable pattern designs. **f**) Light intensity measurements for various system configurations. **g)** Temperature measurements over three cycles of 10 min UV exposure with 3 min intervals. **h)** Light intensity measurements over three cycles of 10 min UV exposure with 3 min intervals.

**Supplementary Figure 2.**
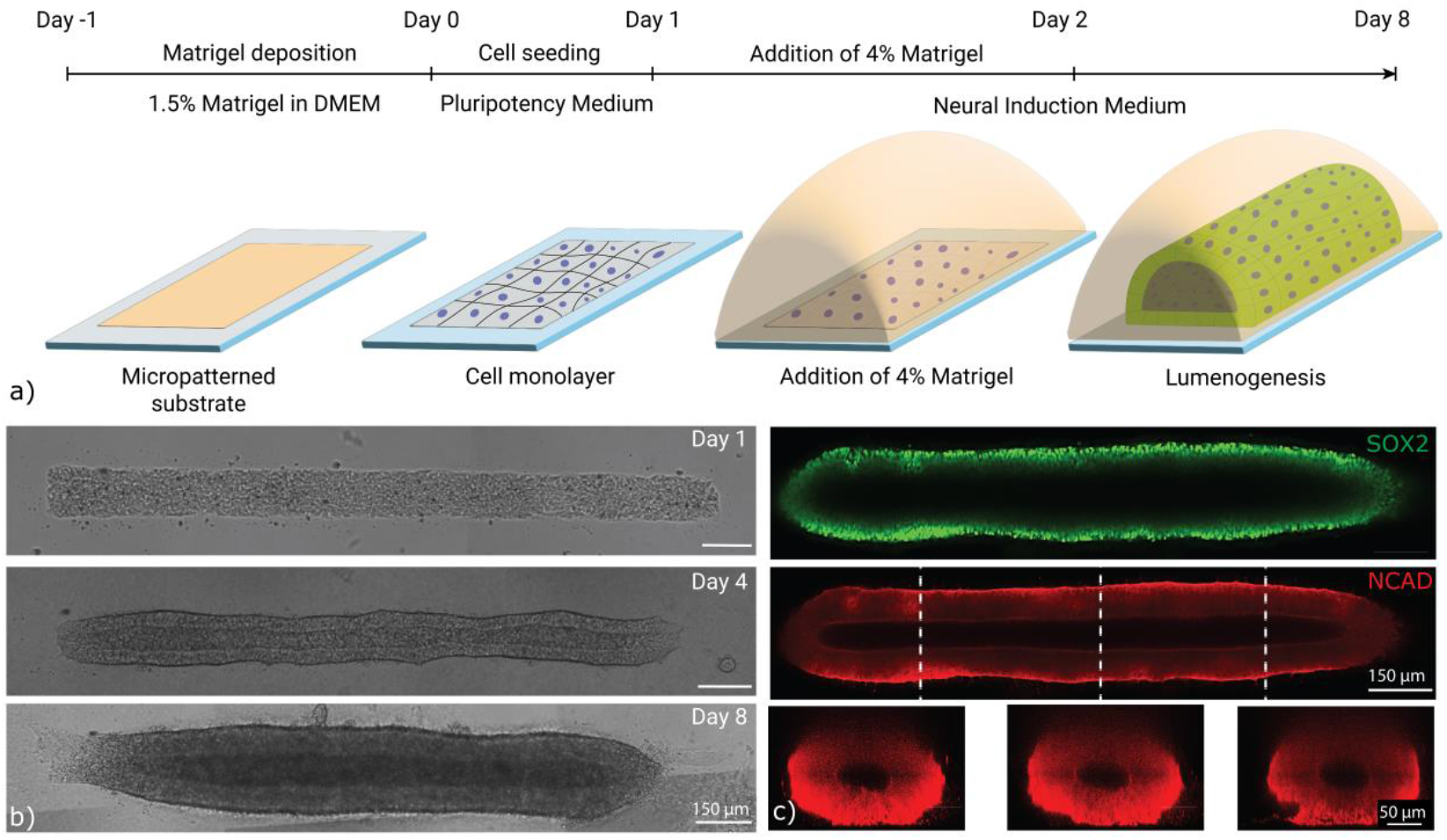
a) Experimental scheme showing the steps to obtain hPSC-derived neural organoids with single-lumen. b) Representative images of organoid development over 8 days of culture. c) Day 8 single-lumen neural organoids obtained starting from patterns generated by the SCAMPI device.

**Supplementary Figure 3.**
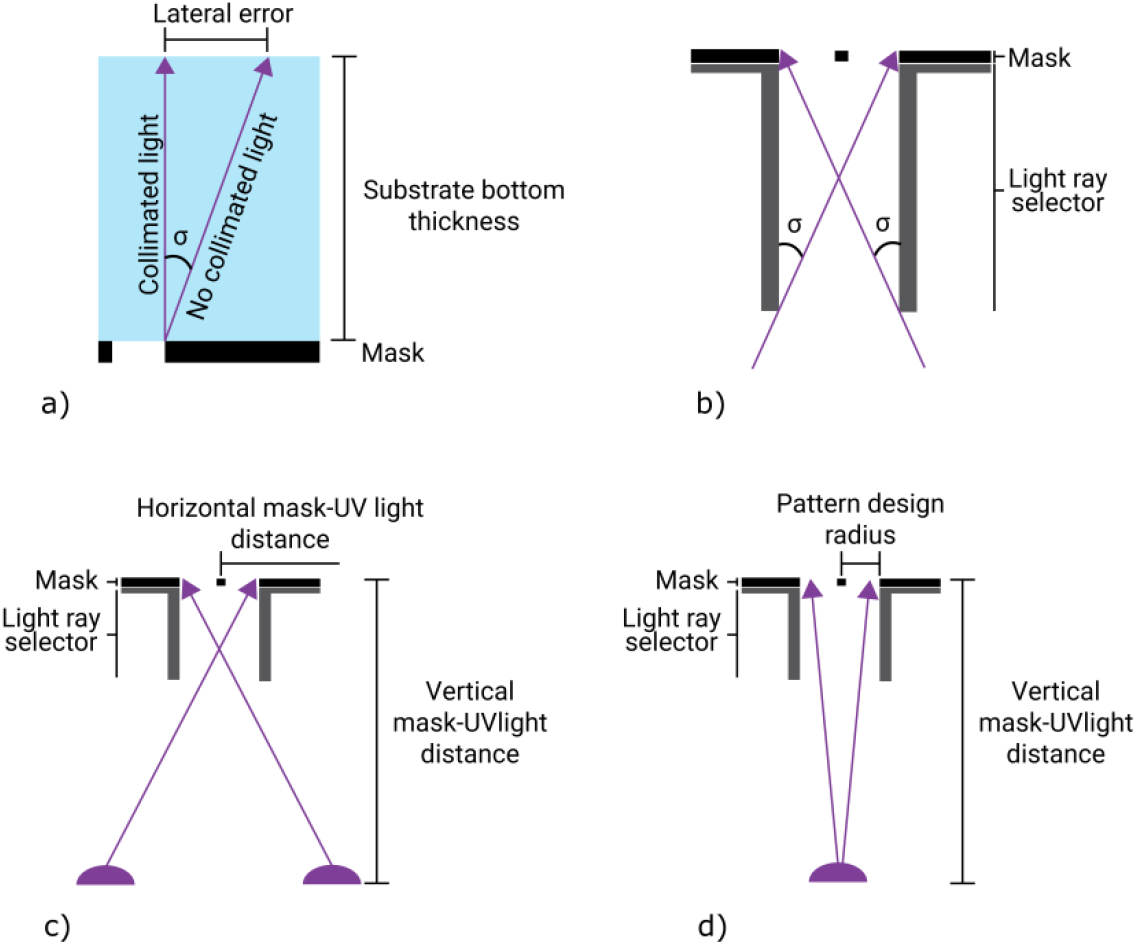
a) Schematic of light ray angle calculation to achieve a lateral error below 15 µm. b) Schematic of the light ray selector showing calculation of depth required to prevent light rays with angles exceeding 3.37° from passing through the mask. c) Schematic for calculating mask distance from the UV light source for the 96-well plate configuration. d) Schematic for calculating mask distance from the UV light source for the glass culture dish configuration

